# Beyond Viral Load: A Mechanistic Model of Virus–Immune Dynamics and Age-Dependent Influenza Severity

**DOI:** 10.64898/2026.03.26.714633

**Authors:** Reagan R. Johnson, Havilah Neujahr, Rodolfo Blanco, Esteban A. Hernandez-Vargas

## Abstract

Influenza infection results from tightly coupled interactions between viral replication, host immune responses, and the emergence of clinical symptoms. While mathematical models have extensively characterized viral and immune dynamics, the mechanistic link between immune activity and disease severity remains poorly understood.

Here, we develop an integrative, within-host modeling framework that explicitly connects infection dynamics, immune responses, and symptom manifestation through a unified dynamical system. Using murine influenza data, we incorporate key immune components alongside a mechanistic representation of symptom progression, quantified via host weight loss. Our analysis identifies inflammatory signaling, particularly TNF-***α***–mediated pathways, as a central driver linking immune activity to symptom severity. Importantly, we demonstrate that age-dependent alterations in immune regulation reshape this coupling: aged hosts exhibit prolonged inflammatory responses that amplify and sustain symptom burden despite comparable viral kinetics.

These results highlight that disease severity cannot be inferred from viral load alone, but instead emerges from the dynamical interplay between immune regulation and host physiology. This framework provides a quantitative basis for understanding age-specific morbidity and offers a foundation for designing interventions that target immune-mediated pathology rather than viral replication alone.

## 1 Introduction

Influenza A virus (IAV) is a negative sense RNA virus that manifests as a highly contagious respiratory infection [1]. In most instances, the infection is mild and easily cleared by the host without intervention. However, for individuals at high risk for virally related complications, an IAV infection can be life threatening. The severity of an influenza infection is largely driven by host factors, such as pre-existing immunity, age, underlying medical conditions, and genetic susceptibility. These factors impact the ability of the host’s immune system to properly respond to infection [2]. A dysregulated immune response can lead to detrimental consequences through excess inflammatory cytokine production, delayed responsiveness, and an inability to effectively clear infection [3].

There is a large body of research modeling the immune response to influenza [4–9]. However, less work has been done in modeling an explicit connection between the immune response and disease severity, where, often, it is no considered when modeling in-vivo infections. To address this gap, we develop a mechanistic within-host modeling framework that explicitly links viral dynamics, immune responses, and disease severity within a unified system. Building on established models of influenza infection, we incorporate key innate and adaptive immune components, as well as introduce a minimal, yet interpretable, representation of symptom progression through host weight dynamics. This approach allows us to move beyond treating morbidity as a secondary or empirical outcome and instead model it as an emergent consequence of immune activity.

Disease severity observed during experiments using murine models is measured using several metrics, one of which is often some variation of scoring the severity of sickness behavior. This scoring depends on the observations made by the experimentalist. While this metric is useful for comparisons between treatment groups within the experiment, as well as determining humane endpoints, it is, by nature, qualitative [10]. Weight over the course of infection however, is a quantitative measurement of disease severity, making it more apt for modeling. As such, our model makes use of weight as the measure of disease severity. Previous work in modeling weight loss due to viral infections has been published. Myers et al. [11] identified a non-linear relationship between percent weight loss and percent of lung lesioned or inflamed. Pinky et al. [12] found an alignment in the impact of the cumulative area under the curve of predicted infected cells and percent weight loss between different viral infections. Our work differs in that we investigate how inflammation may be inducing this loss of weight during infection.

The immune response’s impact on weight involves a complex network of cytokines impacting feeding behavior, as well as increasing nutrient requirements of immune cells [13]. In this model, we focus only on the former. Sickness-associated anorexia (SAA) is a shared trait among organisms where infection induces reduced caloric intake [14]. Several cytokines, such as IL-8, IL-6 and TNF-*α*, are known to induce anorexia [15]. Of interest to us is TNF-*α*, which has been shown to induce the secretion of leptin by adipocytes [16]. Leptin is a metabolic hormone that regulates food intake by suppressing appetite. It has been found that TNF-*α* has a strong positive correlation with leptin in patients who are septic or experiencing systemic inflammatory response syndrome (SIRS) [17]. Additionally, it has been reported that depletion of TNF-*α* in mice infected with influenza reduces weight loss and disease severity [18].

One group of individuals that is considered high risk for severe influenza infections is individuals 65 years of age or older. The CDC estimated that in 2024-2025, 57.4% of IAV-related hospitalizations and 70.6% of IAV-related deaths in the U.S. were patients 65 years of age and older [19]. This increased susceptibility is largely attributed to age-related changes in the immune system, such as a decline in the ability to clear pathogens, reduced chemotaxis and phagocytosis by innate immune cells, decreased interferon production, as well as reduced levels and delayed responsiveness of T cells [20, 21]. Aged individuals also exhibit an imbalance of inflammatory cytokines, which includes a higher baseline of inflammation, as well as a delayed increase and clearance of these cytokines during infection, resulting in prolonged symptoms and heightened risk of inflammation-related complications [22].

Importantly, age-associated, low-grade increases in the circulation of TNF-*α*, along with other inflammatory cytokines, were found to be strong predictors of all-cause mortality risk in elderly individuals [23]. This chronic, low-grade inflammation is also believed to contribute to the onset of a cytokine storm during infection, which is associated with severe disease pathology [24]. As such, in this work we focus on an aged immune response as a model for severe disease.

The impact of age on disease severity is not unique to humans, with similar trends also being observed in mice. Toapanta [25] reported that aged mice given a non-lethal dose of H1N1 Influenza had higher morbidity and prolonged viral clearance compared to young-adult mice due to age related impairments of the immune response. Using data from young and aged mice, we investigate how age-related changes in immune regulation reshape the coupling between inflammation and symptom severity.

## 2 Methods

### Modeling Infection and Immunity

To investigate the impact of aging on immunity, we propose a model that builds from the the work published by Price et al. [6], which we adapted to the mechanisms of interest, then extended to consider host weight loss (Figure 1). Our model is as follows:

**Fig. 1.**
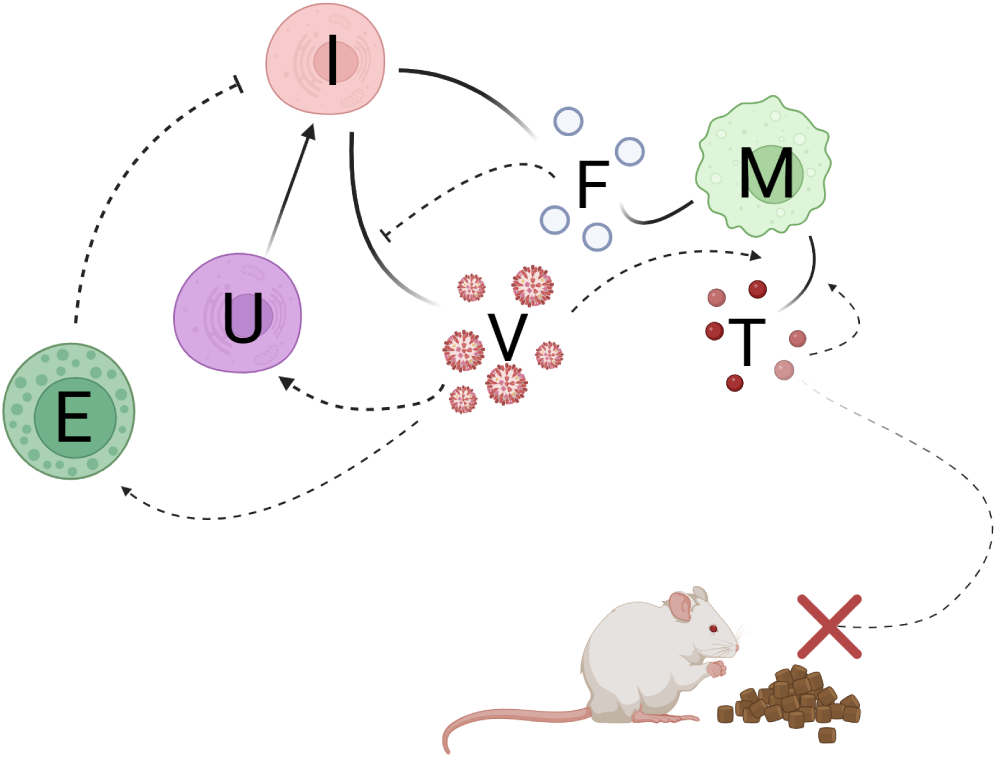
Visual scheme of immune response and disease severity model. Uninfected cells (*U*) are infected by virus (*V*). The virally infected cells (*I*) then produce additional virus and IFN-I (*F*), as well as recruit macrophages (*M*). CD8^+^ T cells (*E*) proliferate in response to virus and kill infected cells. Macrophages also produce IFN-I, which inhibits viral production, and TNF-*α*, which, along with virus, promotes additional TNF-*α* production. The presence of TNF-*α* reduces nutrient intake by the infected mice. Created in BioRender. Johnson, R. (2026) https://BioRender.com/xqchh05

#### Uninfected cells (*U*)

Virus (*V*) infects uninfected cells at rate *β*, upon which they move directly to an infected, virus producing state.

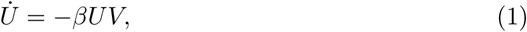

#### Infected cells (*I*)

Once infected, cells will die one of two ways. One, they die by reaching the end of their infected life cycle (*δ_I_*_1_). For Influenza infected cells this is estimated to be around 12 hours [4, 26]. Thus we fix the rate as *δ_I_*_1_ = 2. Two, infected cells die via CD8+ T cell (*E*) mediated-killing, where infected cells are killed at rate *δ_I_*_2_.

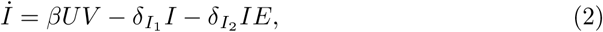

#### Influenza virus (*V*)

Additional virus is produced by infected cells at rate *p*, but the presence of Type I interferon (*F*) reduces this production in a concentration dependent manner, as assumed by Price et al. [6]. The strength of this effect is mediated by the constant *k_F_*. Virus is then cleared from the lungs at a constant rate *c*.

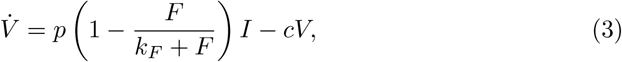

#### Macrophages (*M*)

The initial population of macrophages is represented by *S_M_*, which we derive from the state condition where *V* = *I* = 0 such that *S_M_* = *δ_M_ M*_0_. Infected cells recruit additional macrophages to respond to infection via a hill function. Macrophages are assumed to clear from the lungs at a constant rate *δ_M_*.

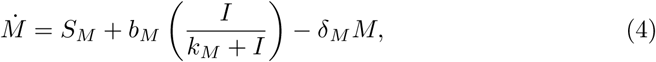

#### CD8^+^ T cells (*E*)

CD8^+^ T cells are a component of the adaptive immune response. Their activation and action have been well characterized in previously published models [11, 27], including in models studying the impact of an aging immune system [28]. We take a simplified approach to modeling T cell activation where we allow virus to directly induce T cell proliferation. Much akin to macrophages, this is a subrogate of a complex process of cellular recruitment and proliferation. This CD8^+^ T cell dynamic was presented in Whipple et al. [28]. Here, *S_E_* is considered to be the proliferation rate during homeostasis, which is derived from the steady state where *V* = *I* = 0 such that *S_E_* = *δ_E_E*_0_, *r* is the CD8^+^ T cell proliferation rate in response to virus, and *k_E_* is the half saturation constant for viral activation of T cells.

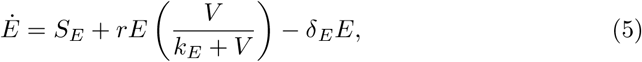

#### Interferon I (*F*)

Type I interferons are antiviral cytokines produced by several different cell types in response to infection. Type I IFN stimulates the activation of antiviral genes. These genes are capable of interfering with various stages of the viral life cycle [29]. Type I IFN is primarily produced by infected cells, which is reflected in our model as rate *q_F_*_1_. Our model also considers macrophages’ production of IFN-I at rate *q_F_*_2_ [30]. The half life of Type I IFN is denoted by *δ_F_*.

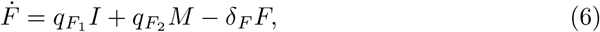

#### TNF-*α* (*T*)

Our model considers macrophages’ production of TNF-*α* in response to both virus and TNF-*α* (*b_T_*_1_) [31]. The strength of viral signaling on TNF-*α* production is taken to be a hill-like function with the constant *a*. We also assume there is an additional rate of TNF-*α* production by macrophages (*b_T_*_2_), which was estimated from TNF-*α* and macrophages on day 0. This was implemented to better follow the observed trends of TNF-*α* (Figure 5). The half-life of TNF-*α* is represented by *δ_T_*.

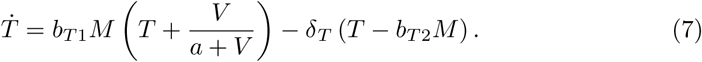

### Modeling Disease Severity

In order to construct our model for disease severity, we first needed a baseline weight model. As these mice are in captivity, we know they are offered a consistent amount of food each day. As such, we assume that under healthy conditions, they are consuming a constant amount of nutrients each day, which we denote by *eW*_0_, where *W*_0_ is the initial weight of the mice. This mechanism allows for bounded growth, which was observed in non-infected mice over the course of the experiment Toapanta [25]. By similar logic, we assume that the mice have a constant rate of energy use proportionate to their body weight. This is represented by the second term in our weight equation, −*δ_W_*, which broadly represents the energy needed for regular metabolic processes. Thus, in non-infection conditions, the modeled weight dynamic is as follows:

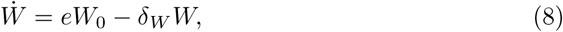

Our model assumes that TNF-*α* has an immediate impact on nutrient intake in a concentration dependent manner. This assumption comes from experimental observations of TNF-*α* increasing the circulation of the hunger-suppressing adipokine leptin in a dose dependent manner [32]. As such, in this model, an excessive concentration of TNF-*α* results in near complete cessation of eating while low concentrations will have little to no impact. All together, our disease severity model is as follows:

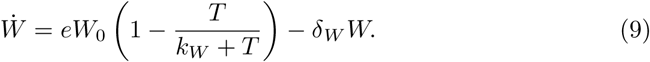

### Experimental Data

The data used in our modeling is a subset of a much larger collection of data that has been thoroughly described in its original publication [25]. Briefly, female BALB/c mice, aged 12-16 weeks (young) and 72-76 weeks (aged), were intranasally inoculated with 50-100 PFU of a mouse-adapted strain of influenza virus, A/Puerto Rico/8/34 (PR8) (H1N1). Mice were then monitored for morbidity for 19 days. Weight and symptom scores were recorded each of the 19 days. Lungs were harvested on days 0, 1, 2, 3, 5, 7, 9, 11, 15, and 19 post-infection. Lungs were then homogenized for data collection. Viral titer was determined by plaque assay, and both ELISA and a multiplex assay were used to measure cytokine and chemokine concentrations. Immune cell populations were characterized using flow cytometry [25]. Of the data collected in the original experiment, our model uses the measurements of viral titers (PFU), IFN-I (*α* and *β*) (pg/ml), TNF-*α* (pg/ml), interstitial macrophages (cell counts per gram), and CD8^+^ T cells (cell counts per gram) [25].

In addition to immune response data, we also use the weights of the infected mice, which are recorded in percent weight with respect to the mouse weight on day 0 of infection. Of the young mice weight dataset, one mouse from the longitudinal data was excluded from the model since its lack of weight loss indicated it likely did not receive a full dose of PR8.

### Parameter Fitting

Parameter fitting was done using the global optimizer differential evolution provided by the python library Scipy [33]. This optimizer minimizes the cumulative cost function of each data set since the data sets used had differing magnitudes. Using a cumulative, non-normalized cost function caused the global optimizer to overly prioritize the data sets with large magnitudes, such as viral titers, and neglect the smaller in magnitude data sets, such as TNF-*α*. Thus, we used a normalized RMSE before calculating the cumulative cost function as NRMSE is a non-dimensional metric that allows for comparison between RMSEs of different units [34]. For each of the variables fit to experimental data, the RMSE was calculated as:

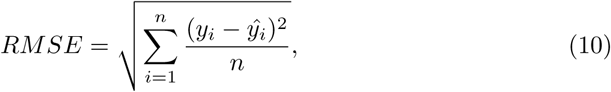

where *y_i_* is the recorded data points and *y*^*_i_* is the modeled value at that time point. Each RMSE is then normalized but dividing it by the difference of the maximum and minimum value within the corresponding data set.

### (NRMSE)_j_

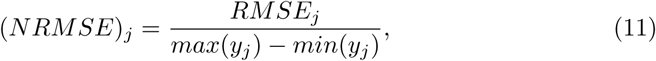

The normalized RMSEs are then summed and the model trajectory with the lowest summed value is considered the model with the best fit.

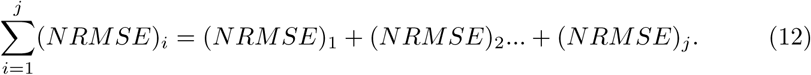

Overall, this improved the performance of the global optimizer, enabling it to produce satisfactory fits. A further constraint was applied to the fitting algorithm in which parameter sets with viral levels exceeding the time of clearance were penalized. To ease computation and fitting times, the immune response parameters were fit first. Those parameters were then fed into an identical model that had the additional equations which accounted for weight. Those additional equations were then fit to the weight data. We adjusted the PFU on day 0 to be 25, rather than the zero recorded. This is in order to align with the value of *V*_0_ in our model. We selected 25 as the mice were inoculated with 50 PFU and assuming *V*_0_ as 50% of the initial dose is a common practice for addressing the uncertainty of the mice receiving a full dose [28, 35]. Parameter fitting results are presented in Table 1.

**Table 1.**
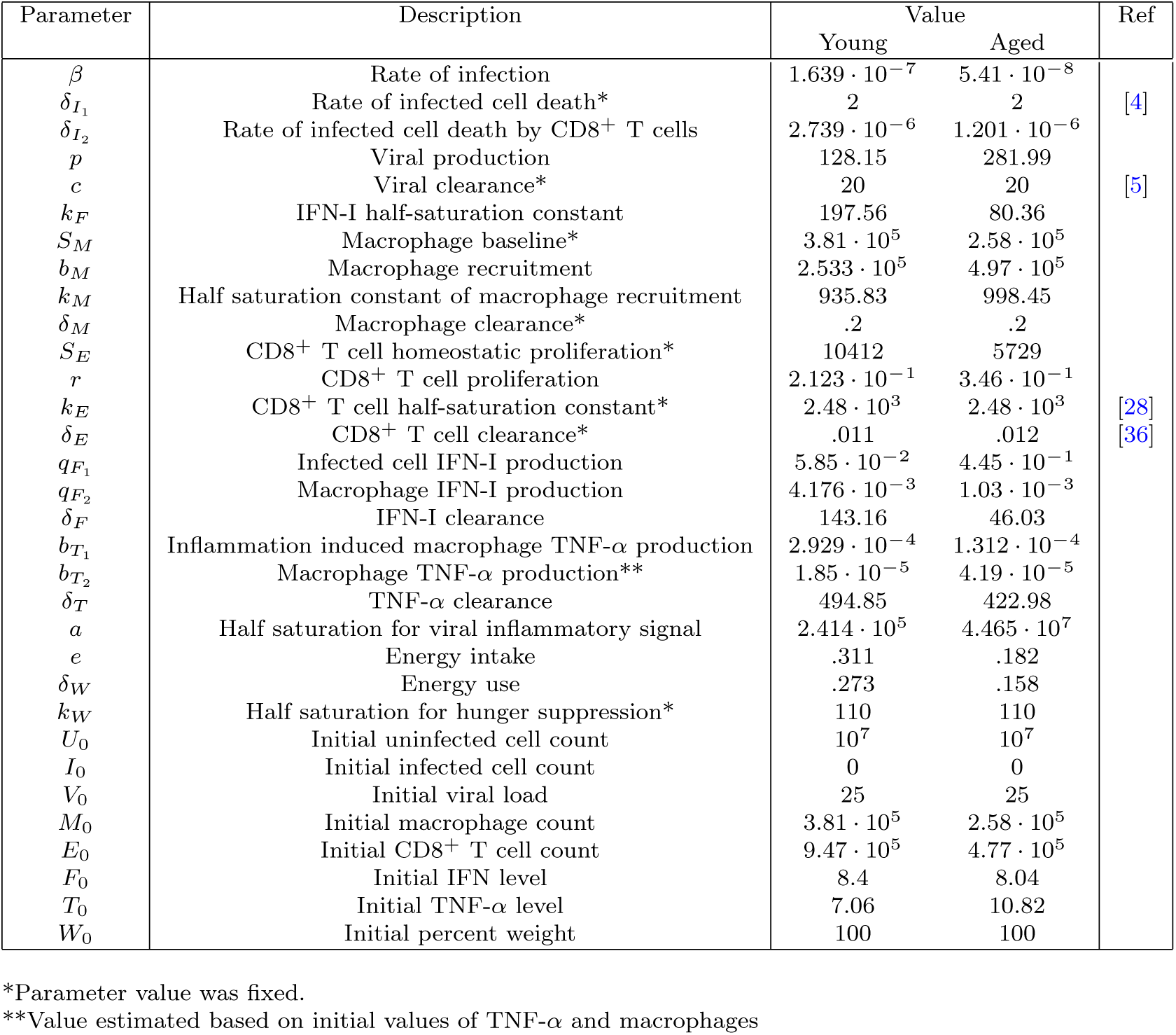
Parameter values of best fit generated from fitting procedure.

**Table 2.**
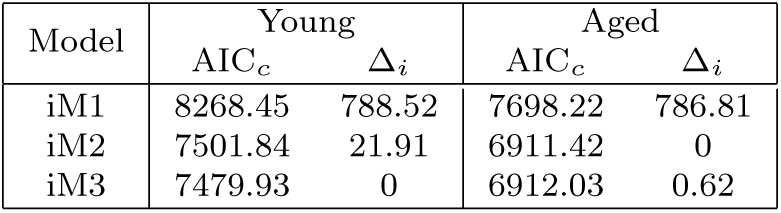
Immune response model selection for both young and aged mice. The Δ*_i_* of iM3 is 0 for the young mice and *<* 1 for the aged mice, which we argue is a small enough difference to consider iM3 to be performing as well as iM2 [37]. As such, this work centers around the results of iM3.

### Model Selection

We considered three models of the immune response to influenza in the development of this work, denoted iM1, iM2, and iM3. Here we focus on iM3 (1)-(9). The first model (iM1) considered an additional variable IL-10, which is an anti-inflammatory cytokine that down-regulates the expression of TNF-*α* by macrophages (see S1 of supplemental material). The second model we considered (iM2) contains an additional term for TNF-*α* promoting additional TNF-*α* production by macrophages (S1 of supplemental material).

Model selection was done using small sample size corrected Akaike Information Criterion (AIC*_c_*), which is a common method of model selection [9, 28]. AIC rewards goodness of fit while penalizing for model complexity to reduce the risk of over-fitting. AIC*_c_* is utilized to correct for small sample sizes [37], and the model with the lowest AIC*_c_* is selected. The formula is as follows:

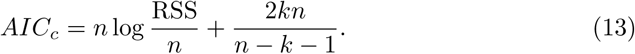

where *n* is the number of data points, RSS is the residual sum of squares of the model’s fit to data, and *k* is the number of free parameters. For ease of interoperability, we present the Δ*_i_* value for the AIC*_c_* of each model for each age [37]:

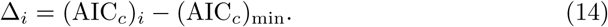

where the model with the lowest AIC*_c_* of each age group will have Δ*_i_* = 0. Models with Δ*_i_* ≤ 2 are also considered to be well supported [37].

Since our model of disease severity uses the immune response model as an input, we calculate the AIC*_c_* using the weight loss data and the parameters pertaining to disease severity separately from the immune response model. Utilizing the immune response model selected using AIC*_c_* (iM3), we compared our two disease severity models (Table 3). The first disease model we considered (dM1) assumed there was a delay in the impact on nutrient intake by TNF-*α* (see S1 of supplemental material). The second disease model we considered (dM2), which we present here, assumes there is no delay in impact by TNF-*α*.

**Table 3.** Disease severity model selection, using iM3 as the input for the immune response model.

| Model | Young |  | Aged |  |
| --- | --- | --- | --- | --- |
| | $AIC_c$ | $\Delta_i$ | $AIC_c$ | $\Delta_i$ |
| dM1 | 710.92 | 28.75 | 552.59 | 36.21 |
| dM2 | 682.17 | 0 | 516.38 | 0 |

**Table 4.**
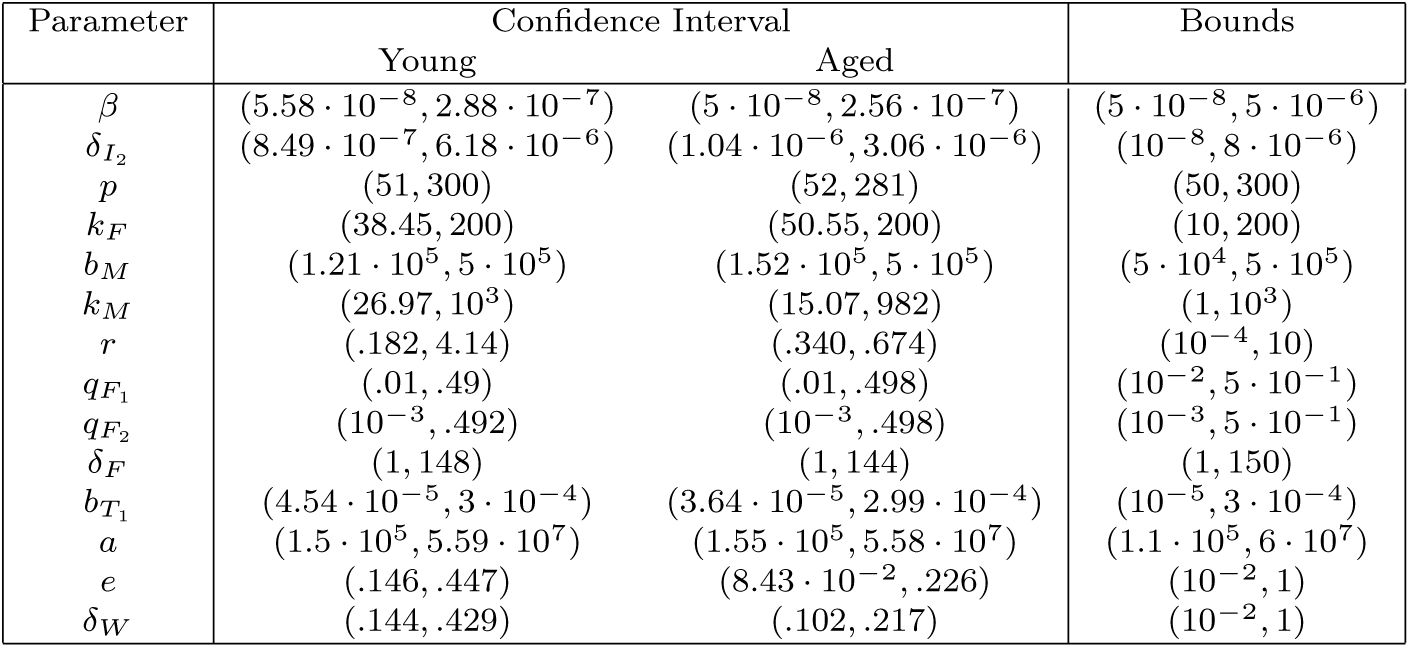
95% Confidence intervals of fitted parameters.

### Parameter Uncertainty

To quantify parameter uncertainty, we utilized nonparametric bootstrap resampling to generate 95% confidence intervals of our fitted parameters. Nonparametric bootstrapping is a common method for obtaining confidence intervals of parameters fit to data [28, 38, 39]. The estimates were constructed by sampling each data set once at each time point. The model was then fit to this same subset of data, and the parameters of best fit were stored. This process was repeated 300 times in order to generate parameter distributions of both age groups. The generated 95% confidence intervals of these distributions are reported in Table 2.

### Sensitivity Analysis

Sensitivity analysis allows us to identify influential parameters within the model, as well as the sensitivity of the model’s behavior to parameter errors [40]. In this work, we take a similar approach as to Whipple et al. [28]. We perform local sensitivity analysis using the One-Factor-At-a-Time (OAT) method, which works by individually varying each parameter while leaving the rest fixed at their value of best fit. We then observe how the behaviors of the responses of interest are impacted by the varied values. The responses we have selected are not only relevant to infection and disease severity, but are also aspects in which we see differing trends between the two age groups. The first of which is time of viral clearance which is taken to be the time point after *t* = 0 in which viral levels fall below the threshold of detection (15) [28].

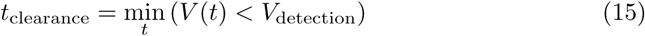

The next two responses analyzed were the peak of TNF-*α* (16) and the time at which that peak occurred. This allows us to observe the sensitivity of the magnitude and timing of the inflammatory response to changes in parameter values.

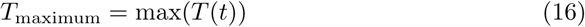

Then, finally, as weight is our measure of disease severity, understanding which parameters are influential to this response is of particular interest. As such, we also analyzed the minimum weight reached (17) and the timing of that minimum.

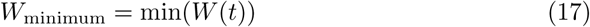

While OAT sensitivity analysis does not consider interactions between factors, it is a commonly used technique as it is easily implemented and interpreted, thus allowing us to observe the local impact that changes in parameter values have on the responses of interest [41]. Here we present a subset of the parameters and responses analyzed, the rest of which are located in *S*2 of supplementary material.

## 3 Results

The model (1)-(9) was able to recapitulate the data of both age groups (Figure 2). The young mice reached higher levels of viral titers, with their peak being 13-fold higher than the aged mice viral peak. Despite this, they cleared the virus quicker than the aged mice. This is reflected by the young mice having a 3-fold higher rate of infection (*β*) and 2.3-fold higher rate of infected cell death by CD8^+^ T cells (*δ_I_*) compared to the aged mice. However, the aged mice infected cells had over a 2-fold higher viral production rate, yet had approximately a 2.5-fold lower threshold for IFN-I-mediated reduction of viral production, implying that IFN-I had a stronger impact in the aged mice. Furthermore, the aged mice had a higher peak of IFN-I, despite the younger mice having a 7-fold higher rate of IFN-I production by infected cells (*q_F_*_1_). The rate of IFN-I clearance was, however, also 3-fold higher for the young mice (Figure 3). This difference in production is consistent with the observations of age related decreases in Type I IFN production [42].

**Fig. 2.**
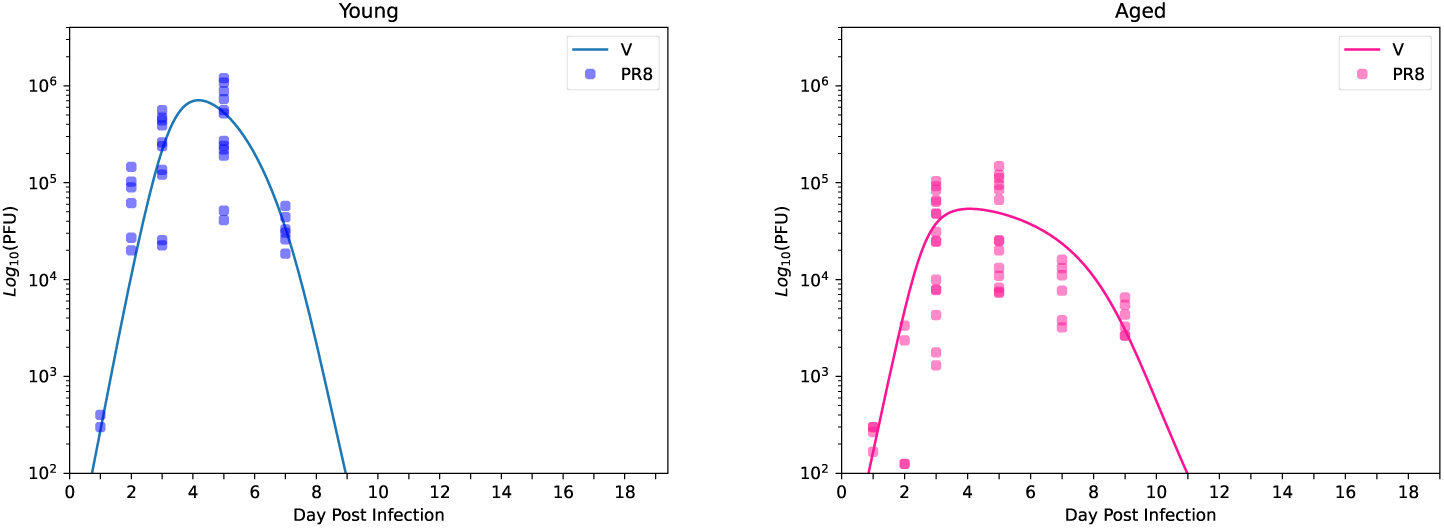
Simulations of the model trajectory for virus (V) for the young mice (left) and aged mice (right)

**Fig. 3.**
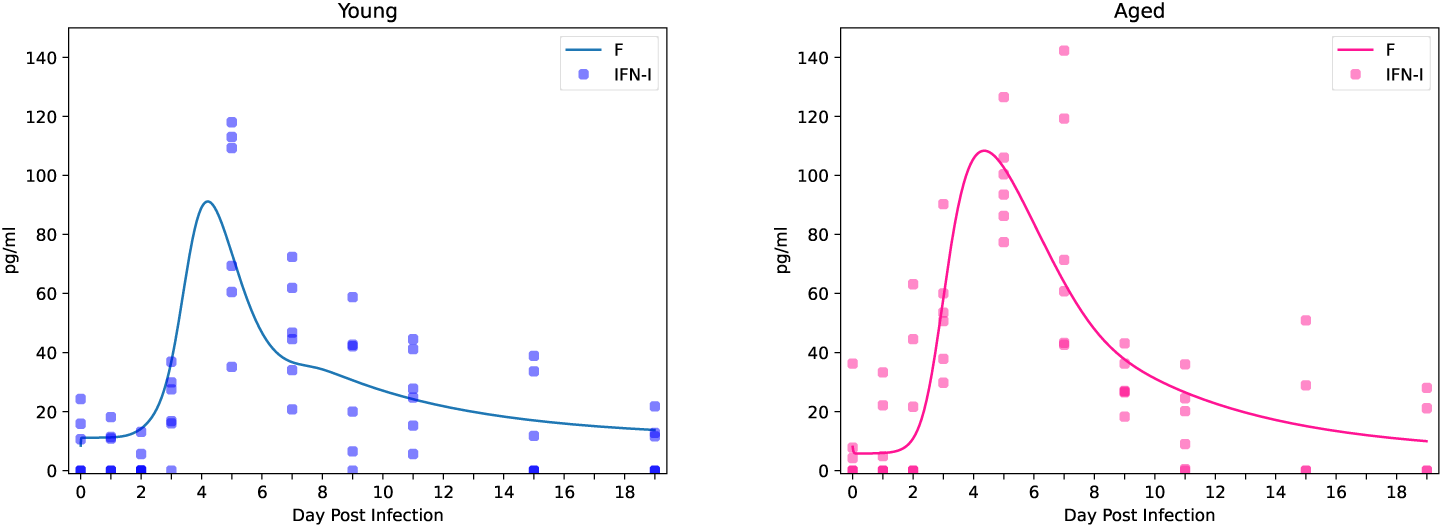
Simulations of the fitted trajectory IFN-I (*α* and *β*) for the young mice (left) and aged mice (right)

Aged macrophage levels were 1.5-fold lower than the young mice on day 0 of infection, but were recruited by infected cells at a 2-fold higher rate (*b_M_*) than the young mice (Figure 4). These recruited macrophages produce IFN-I at rate *q_F_*_2_, of which the young mice had a 4-fold higher rate. Our model was able to follow the increase in macrophage levels for both age groups, but was less apt at following the clearance of the young mice macrophages. This is likely due to the fact that the young macrophage levels on those later days (day 15 and 19) were, for the most part, below the baseline (3.81 · 10^5^), which was estimated from the steady state. Future modeling in which the macrophage baseline is not estimated from the steady state may prove to be more apt at capturing the young macrophage clearance.

**Fig. 4.**
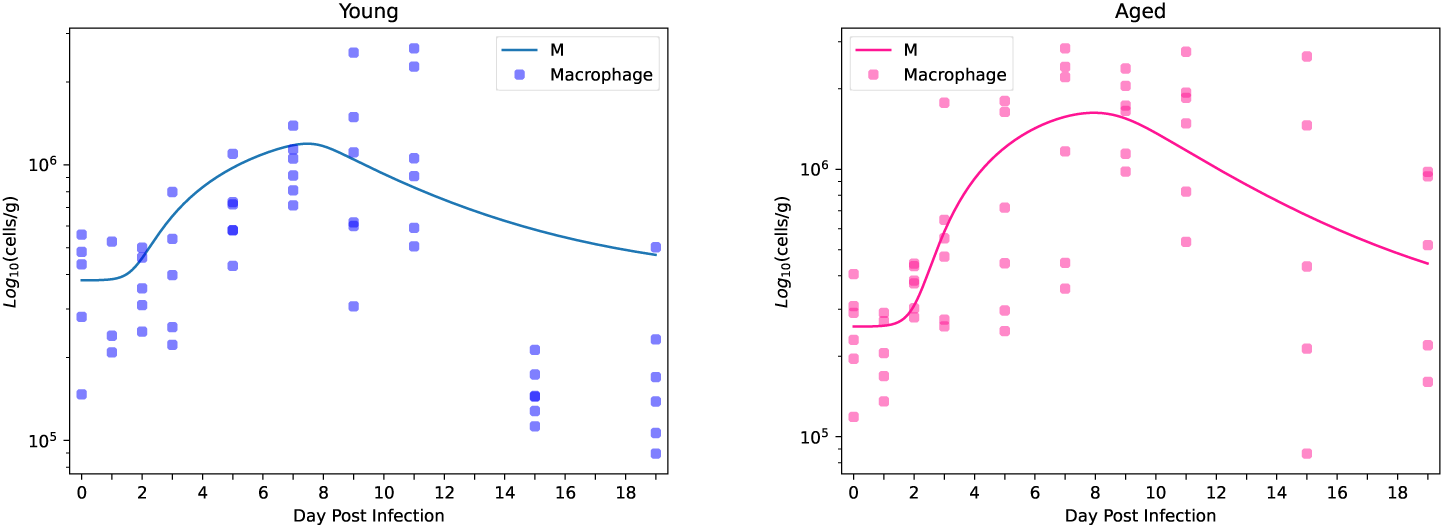
Simulations of macrophages model trajectory for the young mice (left) and aged mice (right)

Macrophages are also responsible for TNF-*α* production. Overall, the two age groups exhibited differing trends in TNF-*α* (Figure 5). The young mice maintained mild levels of TNF-*α*, with its peak coinciding with the peak of macrophage levels (Figure 4). Aged mice on the other hand had a sharper increase and a 1.9-fold higher peak in TNF-*α*. This is, however, not due to a higher rate in inflammation induced TNF-*α* production as the young mice have a 2-fold higher rate (*b_T_* _11_). The aged mice model does exhibit a 2-fold higher rate of baseline TNF-*α* production (*b_T_*_2_), which may be reflective of the observation of chronic low grade inflammation associated with age [43]. This, in combination with a 2-fold higher macrophage recruitment rate (*b_m_*) and lower rate of TNF-*α* clearance, results in a dysregulated inflammatory response observed in the aged mice.

**Fig. 5.**
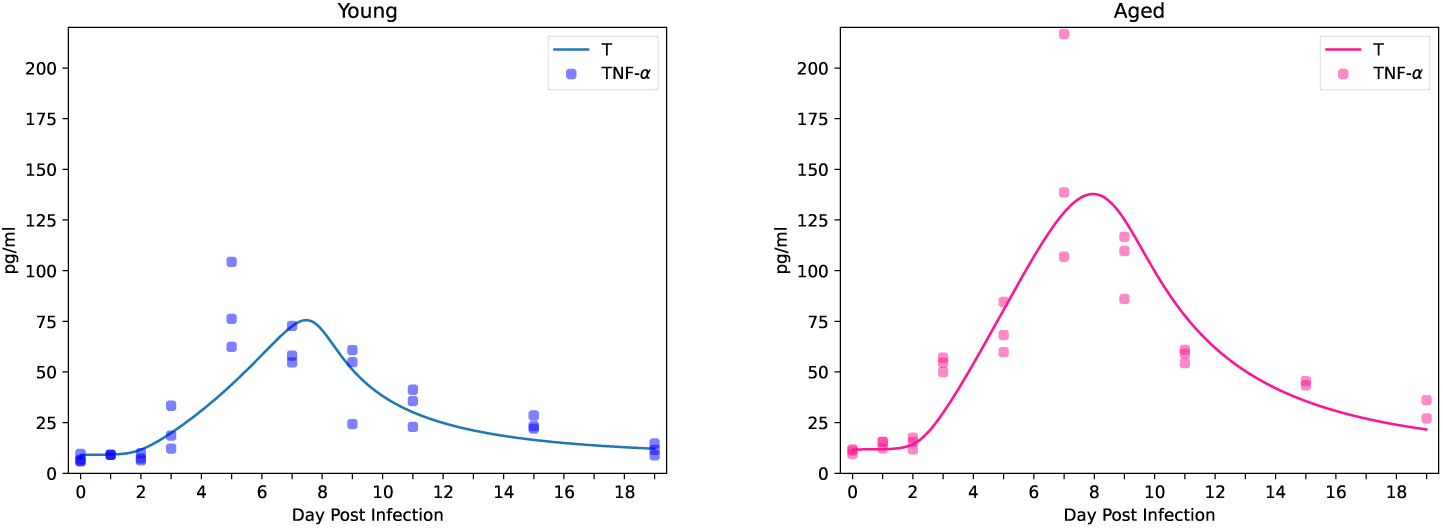
Model trajectory of TNF-*α* for the young mice (left) and aged mice (right)

**Fig. 6.**
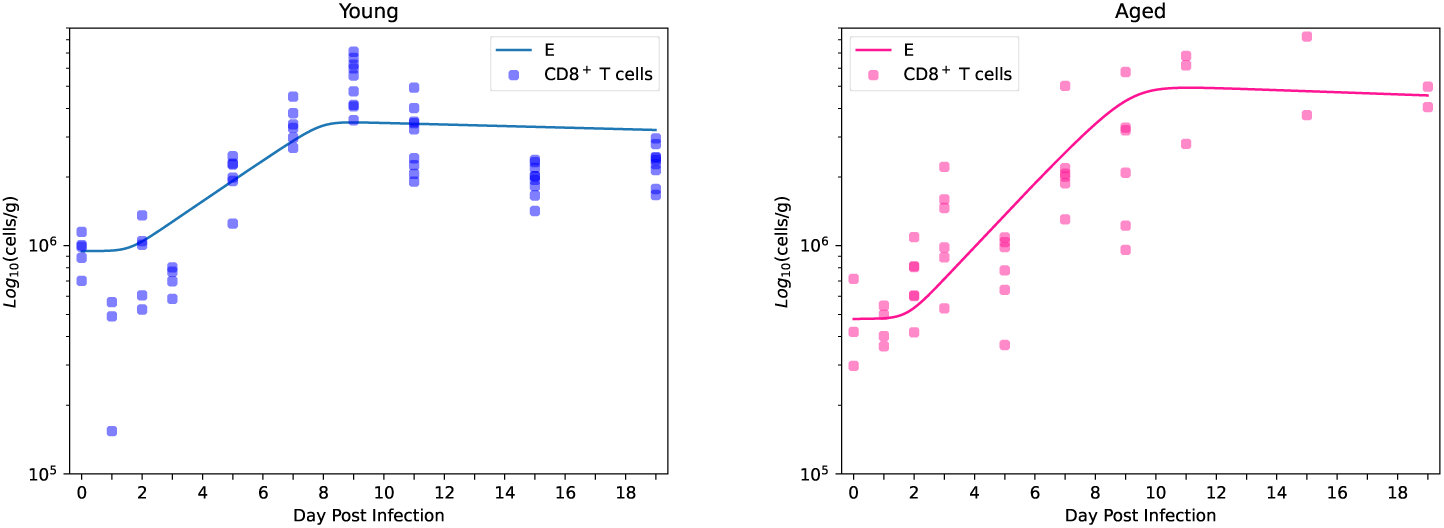
Model trajectory of CD8^+^ T cells for the young mice (left) and aged mice (right)

The young mouse CD8^+^ T cell levels rose then plateaued by day 7 while the aged CD8^+^ T cells continued to rise until day 11. Young CD8^+^ T cells killed infected cells at a 2-fold higher rate, but proliferated slower than the aged CD8^+^ T cells (1.6-fold change). While this a surrogate of a much larger, more complex process, these results do suggest that while aged CD8^+^ T cells are proliferating at a faster rate, they are less effective at clearing infection than their young counter parts.

Weight over the course of infection was a dynamic of interest to us in this model as the goal was to bridge immune response to disease severity. While both groups did lose weight, the young mice recovered the weight lost by the end of experiment. The aged mice on the other hand, lost more weight and did not make the same recovery. The young mice model had higher rates of energy intake and use (1.7-fold change for both), which does align with literature regarding metabolic changes in aging mice [44]. Our model hypothesizes that TNF-*α* is suppressing hunger during infection. To this end, our model was able to follow the general trends in weight loss of the two groups (Figure 7). It was particularly well suited to the aged mice data, which may, in part, be due to the lower amount of variability. It may also suggest that there are additional mechanisms driving the weight loss observed in the young infected mice that were not considered in this model.

**Fig. 7.**
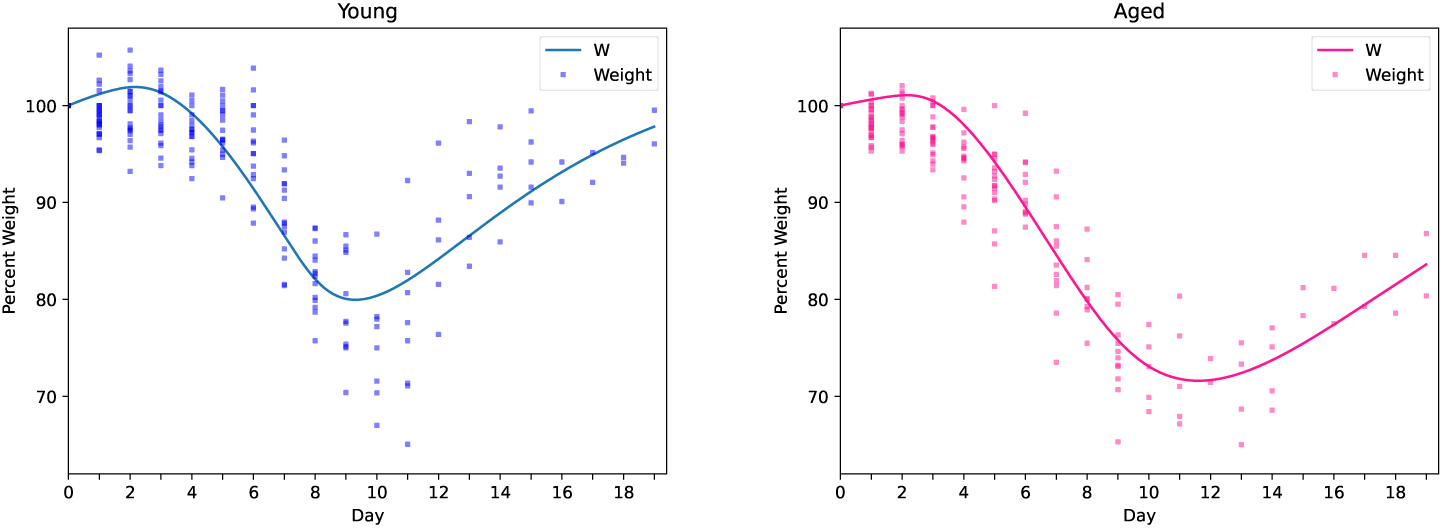
Weight data and trajectory of fit model for young mice (left) and aged mice (right)

### 3.1 Model Analysis

When considering the sensitivity analysis plots in Figure 8, we see that for both the young and aged mice models, an increase in the rate that CD8^+^ T cells kills infected cells (*δ_I_*), as well as increasing rates of CD8^+^ T cell proliferation (*r*), corresponds with a decrease in the peak of inflammation. We see similar trends when looking at the minimum weight reached, which is expected as our model assumes that inflammation and weight loss are tightly connected. These results do align with what we would expect, as higher rates of infected cell clearance result in more efficient clearance of infection, resulting in lower promotion of inflammation.

**Fig. 8.**
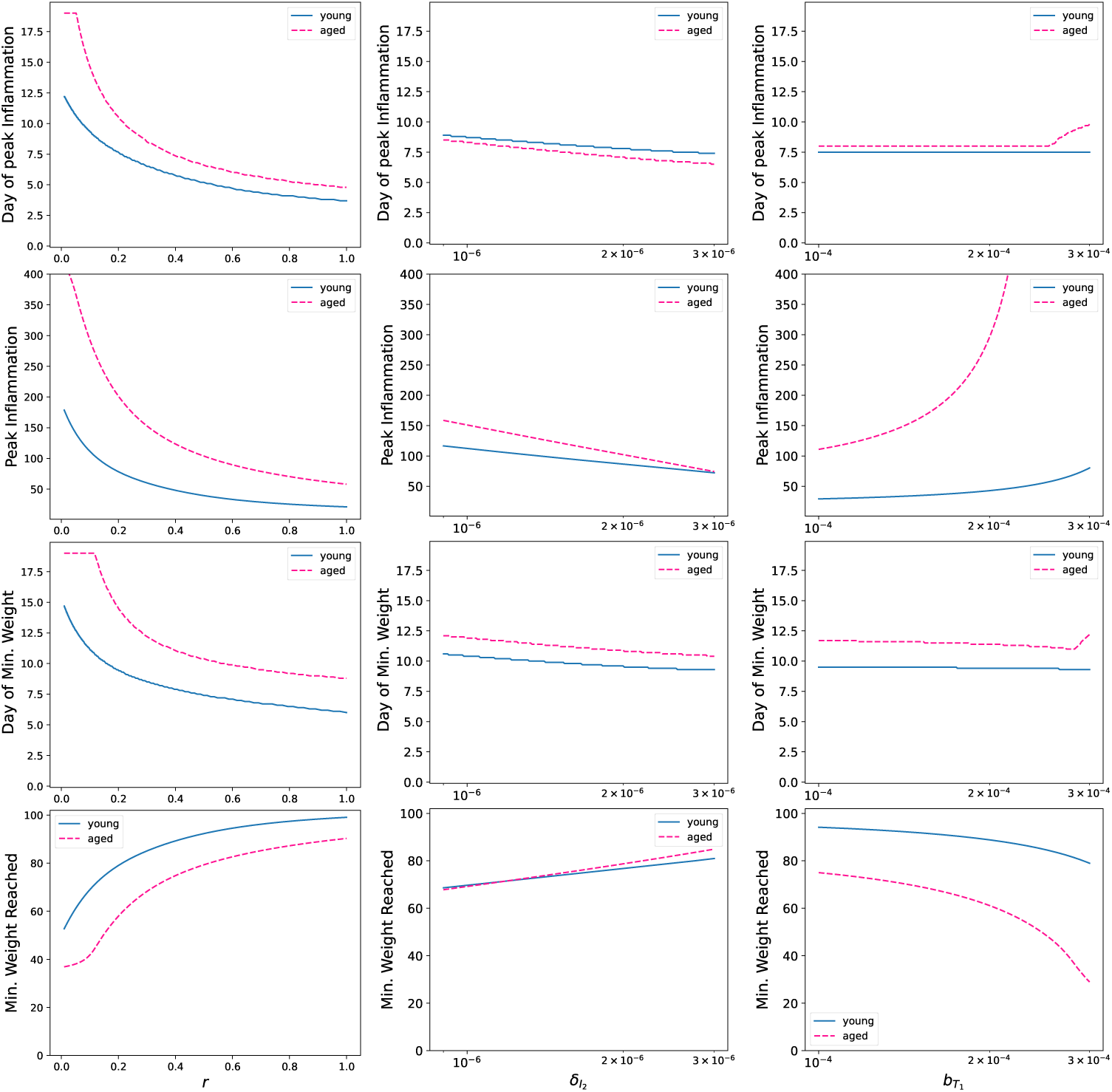
Local Sensitivity analysis results of the parameters *r*, *δ_I_*_2_, and *b_T_*_1_ for both age groups

Turning our attention now to parameters more directly linked to inflammation and weight loss: *b_M_*, *b_T_*_1_, *e*, *δ_w_*, the rate of macrophage recruitment, TNF-*α* production, nutrient intake, and use, respectively (Figures 8 and 9), we see less intuitive results. While for both *b_M_* and *b_T_*_1_ the peak of inflammation rises as the values increase, which does align with what we would anticipate. However, for the aged mice, higher rates of TNF-*α* production results in a much higher peak of inflammation compared to the young mice, well beyond what is observed in the data. The opposite occurs when looking at the rate of macrophage recruitment (*b_M_*), where instead it is the young mice model that exhibits much higher sensitivity to changes at the higher end of the boundary.

**Fig. 9.**
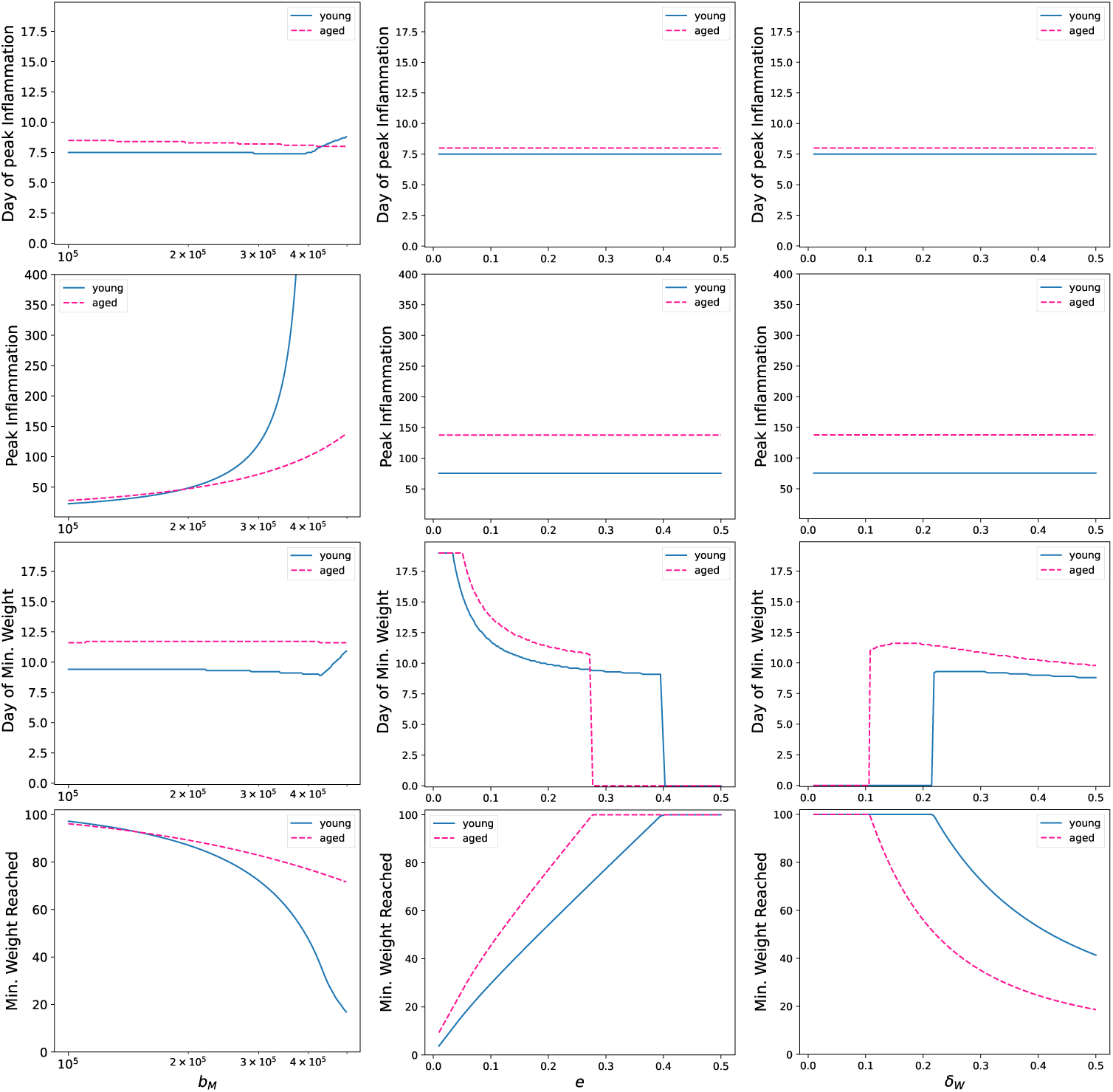
Local Sensitivity analysis results of the parameters *b_M_*, *e* and *d_W_* for both age groups. Plots given a consistent axis for ease of comparison and to capture the behavior of both models

When considering the two parameters that directly correspond to nutrient intake (*e*) and use (*δ_W_*) (Figure 9), we see that the minimum weight reached and the day that minimum occurs appears to be more sensitive to these parameters compared to the others considered. An increase in nutrient intake increases the minimum weight reached approximately linearly until the minimum weight reached is 100% of the initial weight, indicating that weight was not lost during infection. The minimum weight reached is also sensitive to the rate of energy use (*δ_W_*).

In addition to local sensitivity analysis, we also generated 95% confidence intervals using non-parametric bootstrapping. An additional benefit of bootstrapping is it allows for easy identification of correlation between parameters [39]. In Figure 10, we present three pairwise plots of parameter values returned from our bootstrapping procedure. The plots were selected for presentation as they were found to be highly correlated. Correlation was calculated using Spearman rank correlation as this is a nonparametric method which makes no assumptions of the underlying distribution of these values [45].

**Fig. 10.**
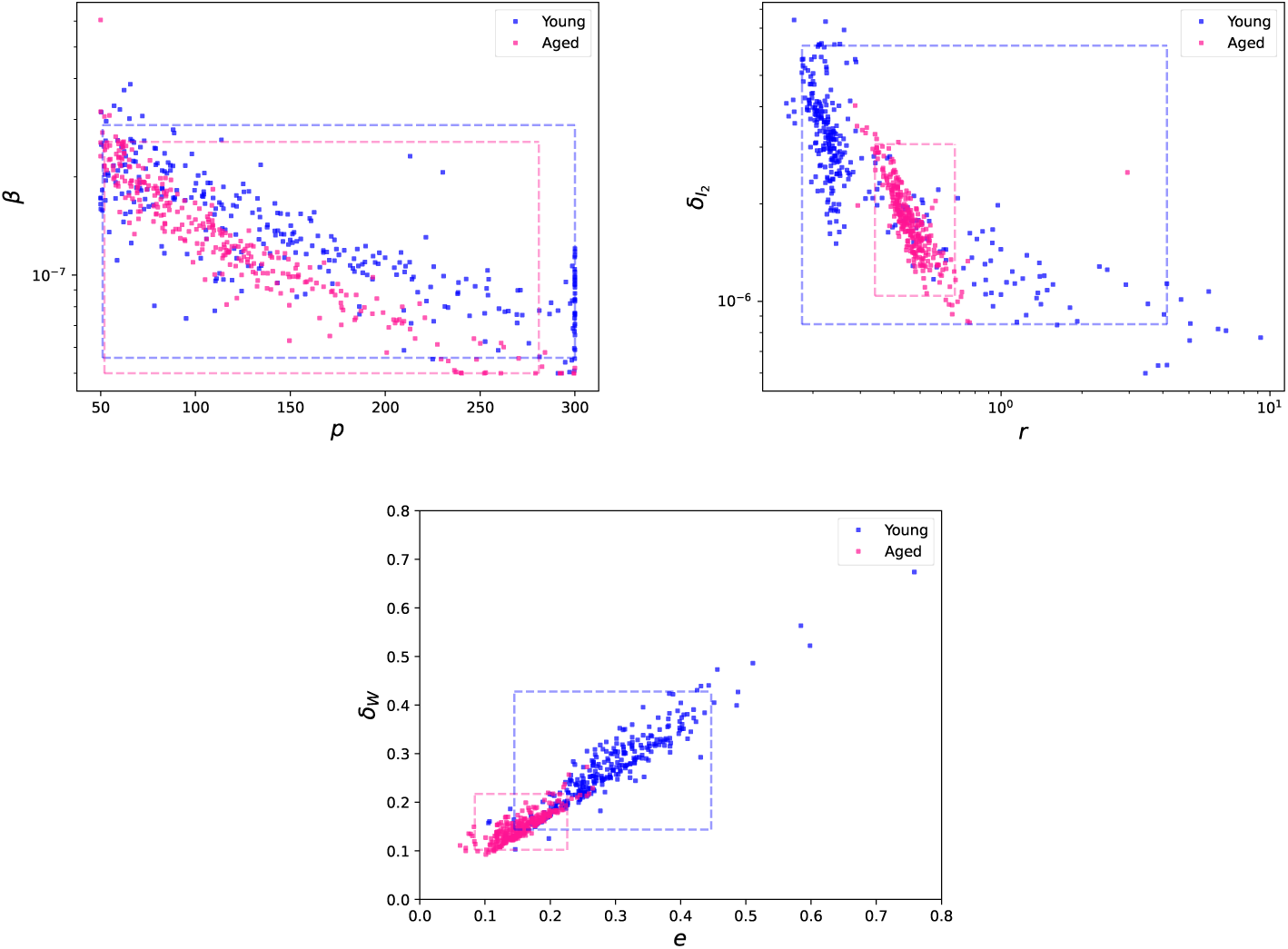
Pairwise plots for values of best fit from bootstrapping procedure. The dashed lines indicate the 95% confidence intervals of the parameters for each respective age group. For both age groups, there is a strong negative correlation between *β* and *p* as well as between *δ_I_*_2_ and *r*. We also observe a strong positive correlation between *e* and *δ_W_* for both age groups

For the parameters *β* and *p*, both models feature a strong negative correlation between the two; −.82 and −.86 for young and aged mice, respectively. This result is not unique to our model. Similar relationships between the rate of infection and viral production have been seen in other within host infection models [46]. Future experimental data that further characterizes the rate of infection and viral replication by infected cells may aid a more unique characterization of these rates.

The other pair of parameters we observe a strong negative correlation between are the rate of CD8^+^ T cell proliferation (*r*) and the rate of infected cell clearance by CD8^+^ T cells (*δ_I_*), which have correlation coefficients −.82 and −.96 for young and aged mice, respectively. This result is reflective of known biological processes. It has been reported that T cells exhibit heterogeneous abilities to clear infected cells and work cooperatively to clear individual target cells [47]. As such, this correlation may be reflective of needing either more, but less-efficient, CD8^+^ T cells or fewer very efficient CD8^+^ T cells in order to clear infection.

The final pair of parameters with a strong correlation is *e* and *δ_W_*, the rates of energy intake and use, respectively. For both age groups, we see a strong positive correlation, .93 and .83 for young and aged mice, respectively. This is not unexpected as energy intake and use balance each other such that high intake and high use may produce the same weight dynamics as low intake and low use [48]. This result is reflective of our model’s limitations due to data availability. If future experiments are able to provide additional information on feeding and activity patterns, a more indepth model could be developed allowing us to better distinguish the intake and use of nutrients [49].

## 4 Discussion

The work presented here proposes a mathematical model of the immune responses influence on disease severity. We applied this model to the data of both young and aged mice non-lethally infected with H1N1 influenza to study how age associated heightened inflammation may be driving the worsened weight loss seen in these aged mice.

This model is derived from the model of the immune response to influenza proposed by Price et al. [6]. Their model featured a large ensemble of immune and inflammatory responses to influenza. In developing our model, we reduced it to the mechanisms relevant to our disease severity model while ensuring it still fit the data. While their larger model may be able to capture a more detailed image of the immune response, our model worked within the limitations of the data available to us and focused on our mechanisms of interest. However, there is still a large number of parameters in this model, a portion of which may not be able to be verified experimentally. In order to make fitting this model feasible, some parameters had to be fixed. These fixed values were, in majority, based on what has been provided in the literature. However, *k_w_* and *d_m_* were fixed qualitatively, and *b_T_*_2_ was estimated from the initial values of TNF-*α* and macrophages.

While our model was able to follow the data, there are limitations. One of which is we only considered macrophages’ cytokine production and not their phagocytosis of infected cells. Our model also considers only the inflammatory phenotype of macrophages. Extending this model to account for macrophage polarization may be a beneficial direction for future work. This may allow one to consider the inflammatory and anti-inflammatory feedback loops that are likely critical underlying mechanisms [50, 51]. Our larger model (iM1) did consider the anti-inflammatory cytokine IL-10 in a manner similar to Price et al. [6], but the added complexity did not further our understanding of how the increased inflammation may be driving weight loss in aged mice.

Previous work has been done in studying a respiratory infection’s impact on weight loss. Pinky et al. [12] observed similarities between the differences of the cumulative area under the curve (CUAC) of their modeled infected cells and the differences of observed weight loss between the treatment groups modeled. They found that groups with more severe weight loss also had a higher CUAC of their modeled infected cells compared to groups with less weight loss. While our model is also focused on connecting infection and disease severity, ours is focused on the immune response and is fit to data. Work has been done connecting weight loss to infection data. Myers et al. [11] found that a saturating function of percent weight loss to inflammation and percent of lung lesioned during infection was well fit to the data and there was a positive correlation between weight loss and inflammation. While our model focuses on how inflammation may be inducing weight loss in IAV-infected mice, the results reported in Myers et al. [11] may support the assumptions made in our model as they identified a relationship between weight loss and inflammation.

The proposed mechanism is, of course, a simplification of the complex relationship between the immune response and disease severity. While TNF-*α* has been connected to weight loss during IAV infections, it is doing so by promoting the secretion of hunger suppressing adipokine leptin [13, 18, 32]. Leptin levels were not available for this dataset, thus we did not consider it directly in our model. However, it has been reported that, following a singular administration of TNF-*α*, leptin levels peaked by 7 hours in mice [32]. This connection has also been found to be clinically relevant in humans. Abd et al. [17] identified a significant positive correlation between leptin and TNF-*α* in patients with systemic inflammatory response syndrome. As such, considering TNF-*α* directly may be an apt approximation for this complex biological process. If future infection experiments are able to provide data on food intake, weight, and leptin levels of the infected mice, a more in depth model could be developed [49].

Our model was able to recapitulate the inflammation and weight trends of both the young and aged mice, suggesting that TNF-*α* may be connected to this loss. Additional factors, such as other inflammatory cytokines like IL-18 or the energetic cost of mounting an immune response, are also likely behind this observation [13]. Models that consider other potential driving factors remain an open opportunity for future study and may provide further insight.

Mice are a common model for the study of human respiratory infections, and aged mice are considered an apt model for elderly human’s immune system [25, 52]. While there are key differences in how influenza manifests in humans and in mice [53], understanding how these age-related changes may be heightening disease severity in mice could help improve our understanding of these changes in humans. This of course remains a pressing issue, as individuals 65 and older are at high risk for influenza infections resulting in hospitalization [54]. In the U.S. this vulnerability is of added concern as our population is aging which furthers the building pressure on our healthcare system. [55]. As such, minimizing severity of infection and risk of requiring intensive care is ever important. Vaccines are the most effective form of infectious disease prevention, and specific formulations of influenza vaccines for aged individuals are available. Despite this, older individuals remain severely impacted by influenza due to age related changes of the immune system [56]. Compounding this issue, influenza vaccination uptake rates have been declining in the U.S. [57].

As a second line of defense, antivirals are available and recommended to be administered as soon as possible to influenza infected patients that are either hospitalized or at high risk for complications [58, 59]; however, antivirals can fail to reduce disease severity. While this failure can be the inability to reduce viral loads due to the virus’ resistance to the drug. The sustained severity is more often due to immune response mediated lung injury caused by an inability to properly regulate the immune response during and after the infection resolves, which is something antivirals can not address [22, 60]. Acute lung injury is characterized by a dysregulated immune and inflammation response which damages lung tissue and can lead to respiratory failure [61], which patients 65 and older have a lower probability of surviving [62]. Treatment targeting an overactive immune system and dampening the pro-inflammatory response along side antivirals may prove to be effective in reducing disease severity during influenza infections [63, 64]. The ideal strategy and duration for such treatment remains an open question for which additional research is necessary, and modeling may be prove to be a useful asset to this process, resulting in improved treatment to minimize the heightened morbidity seen in older patients. Future models that continue to bridge the immune response and symptoms may help further our understanding and identify new directions for research and treatment, thus improving how we care for our most vulnerable community members.

## Declarations

### 4.3 Supplemental Material

A Supplemental Material pdf does accompany this manuscript.

### 4.4 Data Availability

No new data was generated in the creation of this manuscript.

### 4.5 Code Availability

All code and data needed to generate the plots in this manuscript is archived on GitHub.

### 4.6 Ethics approval and consent to participate

Not applicable.

### 4.7 Consent for publication

Not applicable

### 4.8 Author contribution

Conceptualization: Reagan R. Johnson, Esteban A. Hernandez-Vargas; Methodology: Reagan R. Johnson, Rodolfo Blanco, Havilah Neujahr; Writing - original draft: Reagan R. Johnson; Writing - review & editing: Reagan R. Johnson, Esteban A. Hernandez-Vargas, Havilah Neujahr; Supervision: Esteban A. Hernandez-Vargas.

