## Supplementary Information for "Beyond Viral Load: A Mechanistic Model of Virus–Immune Dynamics and Age-Dependent Influenza Severity"

### S1 Alternative Models

#### iM1: Considering the role of IL-10

This model is the closest out of the three to that published by Price et al. [1].

$$\dot{U} = -\beta UV \tag{S1}$$

$$\dot{I} = \beta UV - \delta_{I_1} I - \delta_{I_2} IE \tag{S2}$$

$$\dot{V} = p\left(1 - \frac{F}{k_F + F}\right) I - cV \tag{S3}$$

$$\dot{M} = S_m + b_M\left(\frac{I}{k_M + I}\right) - \delta_M M \tag{S4}$$

$$\dot{E} = S_E + rE\left(\frac{V}{k_E + V}\right) - \delta_E E \tag{S5}$$

$$\dot{F} = q_{F_1} I + q_{F_2} M - \delta_F F \tag{S6}$$

$$\dot{T} = b_{T1} M\left(a_1 T + \frac{V}{a2 + V}\right)\frac{k_T}{k_T + L} - \delta_T\left(T - b_{T2} M\right) \tag{S7}$$

$$\dot{L} = b_{L_1} M\left(a_1 T + \frac{V}{a2 + V}\right)\frac{k_L}{k_L + L} - \delta_L\left(L - b_{L_2}\left(1 - \frac{F}{k_F + F}\right) U\right) \tag{S8}$$

#### iM2: Additional constant influencing inflammatory strength

$$\dot{U} = -\beta UV \tag{S9}$$

$$\dot{I} = \beta UV - \delta_{I_1} I - \delta_{I_2} IE \tag{S10}$$

$$\dot{V} = p\left(1 - \frac{F}{k_F + F}\right) I - cV \tag{S11}$$

$$\dot{M} = S_m + b_M\left(\frac{I}{k_M + I}\right) - \delta_M M \tag{S12}$$

$$\dot{E} = S_E + rE\left(\frac{V}{k_E + V}\right) - \delta_E E \tag{S13}$$

$$\dot{F} = q_{F_1} I + q_{F_2} M - \delta_F F \tag{S14}$$

$$\dot{T} = b_{T1} M\left(a_1 T + \frac{V}{a_2 + V}\right) - \delta_T\left(T - b_{T2} M\right) \tag{S15}$$

#### dM2: TNF-$\alpha$ has a delayed impact on appetite

$$\dot{X} = \frac{lT - X}{\tau} \tag{S16}$$

$$\dot{W} = eW_0(1 - X) - \delta_W \tag{S17}$$

### S2 Sensitivity Analysis

Plots of all local sensitivity analysis performed on model iM3 and dM2 (Figures S1 - S5).

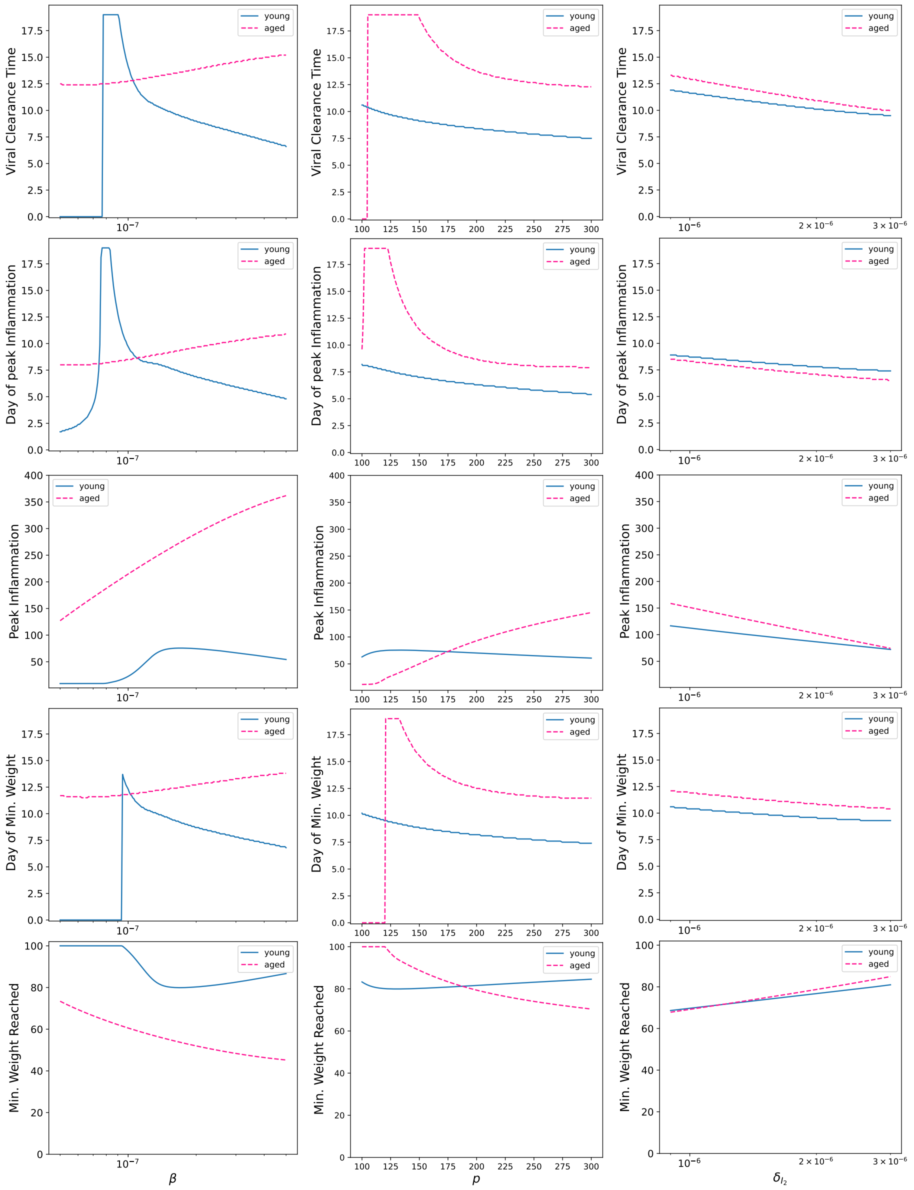


**Fig. S1** Local Sensitivity analysis results of the parameters $\beta$,$p$ and $\delta_{I_2}$ for both age groups

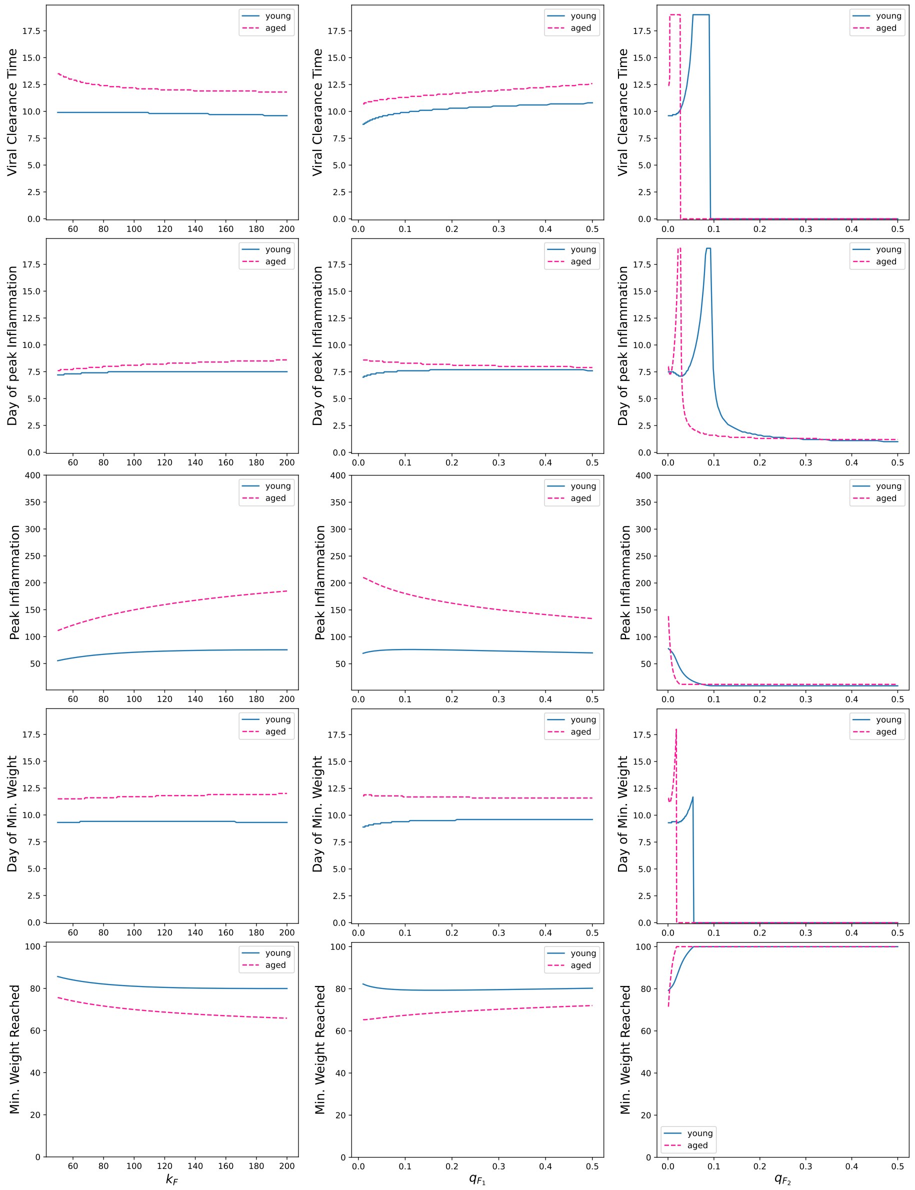


**Fig. S2** Local Sensitivity analysis results of the parameters $k_F$, $q_{F_1}$, and $q_{F_1}$ for both age groups

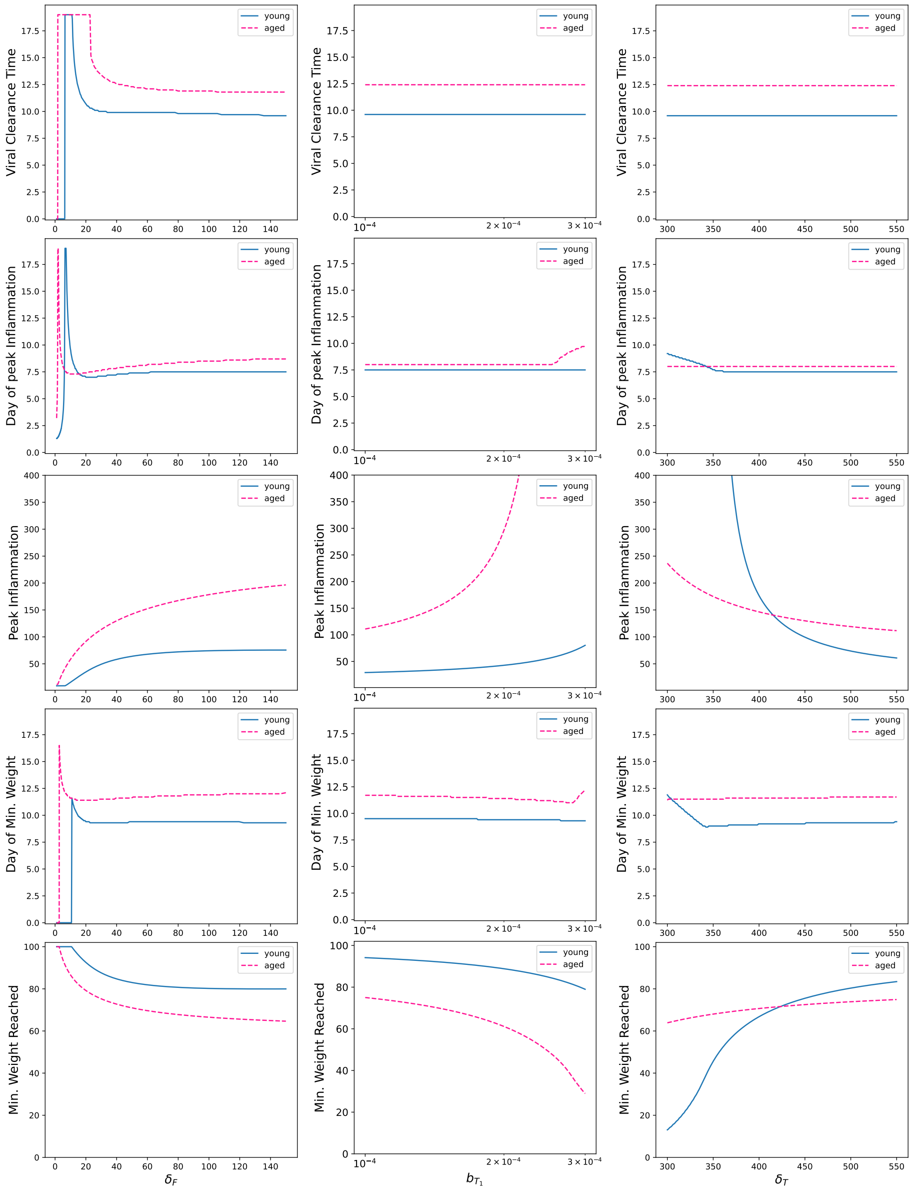


**Fig. S3** Local Sensitivity analysis results of the parameters $\delta_F$, $b_{T_1}$, and $\delta_T$ for both age groups

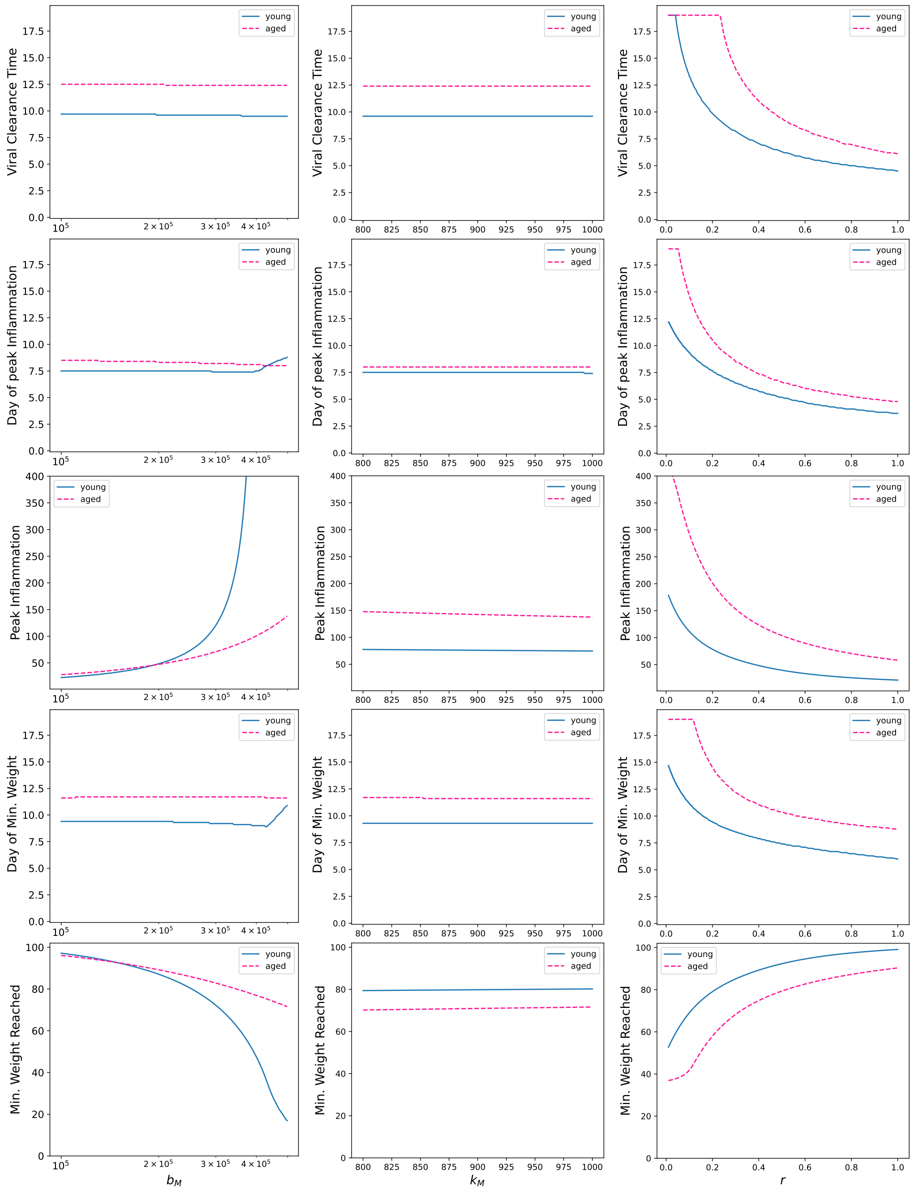


**Fig. S4** Local Sensitivity analysis results of the parameters $b_M$, $k_M$, and $r$ for both age groups

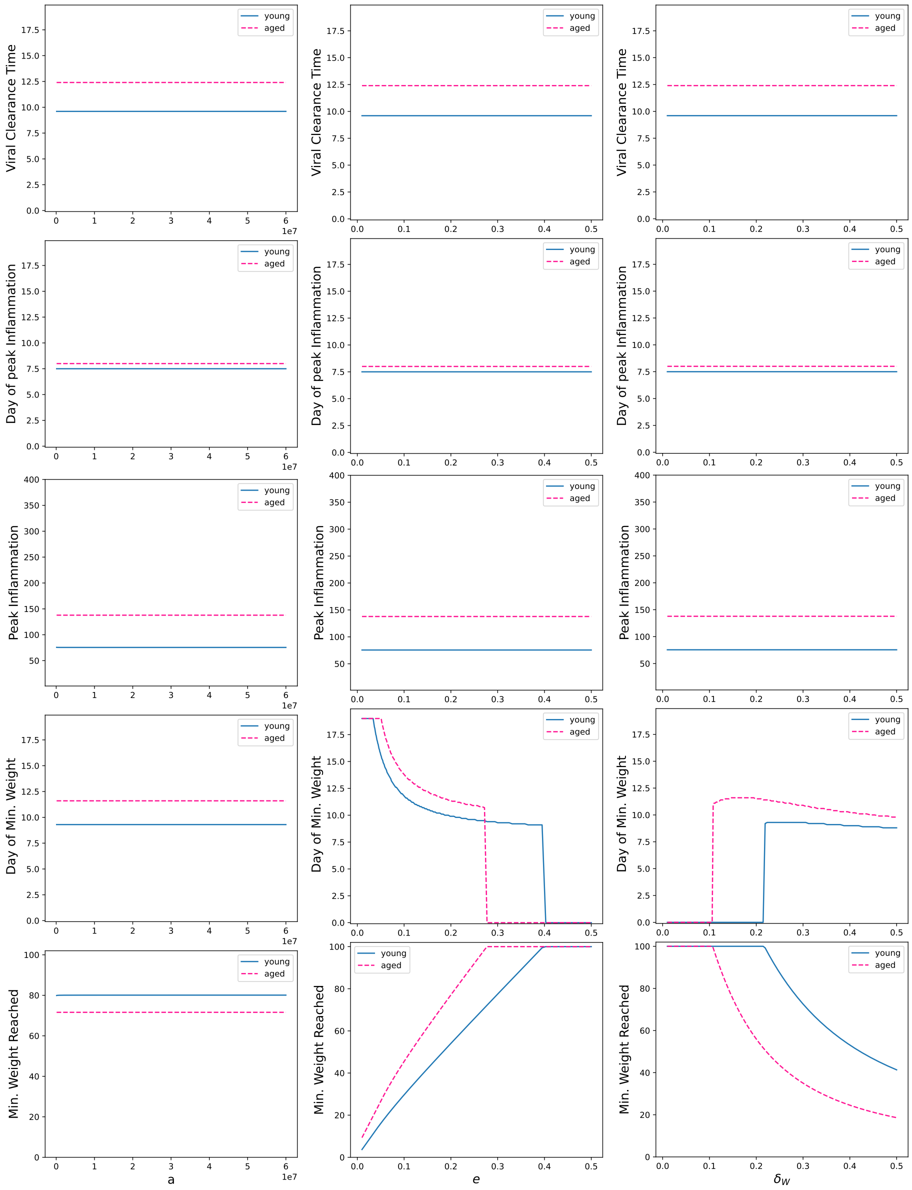


**Fig. S5** Local Sensitivity analysis results of the parameters $a$, $e$ and $\delta_W$ for both age groups

### S3 Bootstrapping

Spearman-Rank correlation of the parameter values of best fit generated from 300 bootstrap samples for both age groups.

| | $k_F$ | $\beta$ | $\delta_{I_2}$ | $p$ | $b_M$ | $k_M$ | $r$ | $q_{F_1}$ | $q_{F_2}$ | $\delta_F$ | $b_{T_1}$ | $\delta_T$ | $a$ | $e$ | $\delta_W$ |
|---|---|---|---|---|---|---|---|---|---|---|---|---|---|---|---|
| $k_F$ | 1.0 | | | | | | | | | | | | | | |
| $\beta$ | 0.029 | 1.0 | | | | | | | | | | | | | |
| $\delta_{I_2}$ | -0.059 | -0.046 | 1.0 | | | | | | | | | | | | |
| $p$ | -0.088 | -0.960 | 0.190 | 1.0 | | | | | | | | | | | |
| $b_M$ | -0.040 | -0.027 | -0.074 | 0.012 | 1.0 | | | | | | | | | | |
| $k_M$ | 0.036 | 0.174 | 0.059 | -0.172 | 0.046 | 1.0 | | | | | | | | | |
| $r$ | 0.038 | 0.191 | -0.863 | -0.338 | 0.097 | -0.063 | 1.0 | | | | | | | | |
| $q_{F_1}$ | 0.201 | 0.373 | -0.141 | -0.420 | 0.030 | 0.169 | 0.238 | 1.0 | | | | | | | |
| $q_{F_2}$ | 0.207 | 0.294 | -0.065 | -0.341 | 0.108 | 0.190 | 0.140 | 0.605 | 1.0 | | | | | | |
| $\delta_F$ | 0.153 | 0.242 | 0.207 | -0.262 | 0.053 | 0.207 | -0.222 | 0.449 | 0.514 | 1.0 | | | | | |
| $b_{T_1}$ | 0.101 | -0.016 | -0.023 | -0.010 | -0.573 | 0.257 | 0.0538 | 0.031 | -0.046 | -0.102 | 1.0 | | | | |
| $\delta_T$ | 0.027 | 0.084 | 0.029 | -0.043 | 0.134 | -0.089 | -0.002 | 0.193 | 0.123 | 0.035 | -0.064 | 1.0 | | | |
| $a$ | -0.008 | -0.063 | -0.042 | 0.054 | -0.018 | -0.004 | 0.084 | -0.031 | 0.025 | -0.052 | 0.025 | -0.068 | 1.0 | | |
| $e$ | 0.092 | -0.0036 | -0.133 | -0.042 | 0.079 | -0.389 | 0.119 | 0.001 | 0.003 | -0.041 | -0.231 | 0.084 | 0.024 | 1.0 | |
| $\delta_W$ | 0.046 | 0.066 | -0.019 | -0.089 | 0.146 | -0.315 | 0.026 | 0.141 | 0.146 | 0.197 | -0.336 | 0.237 | 0.004 | 0.833 | 1.0 |

**Table S1** Pearson Rank Correlation coefficient values of Aged model parameter values generated from bootstrapping.

| | $k_F$ | $\beta$ | $\delta_{I_2}$ | $p$ | $b_M$ | $k_M$ | $r$ | $q_{F_1}$ | $q_{F_2}$ | $\delta_F$ | $b_{T_1}$ | $\delta_T$ | $a$ | $e$ | $\delta_W$ |
|---|---|---|---|---|---|---|---|---|---|---|---|---|---|---|---|
| $k_F$ | 1.0 | | | | | | | | | | | | | | |
| $\beta$ | 0.029 | 1.0 | | | | | | | | | | | | | |
| $\delta_{I_2}$ | -0.059 | -0.046 | 1.0 | | | | | | | | | | | | |
| $p$ | -0.088 | -0.960 | 0.190 | 1.0 | | | | | | | | | | | |
| $b_M$ | -0.040 | -0.027 | -0.074 | 0.012 | 1.0 | | | | | | | | | | |
| $k_M$ | 0.036 | 0.174 | 0.059 | -0.172 | 0.046 | 1.0 | | | | | | | | | |
| $r$ | 0.038 | 0.191 | -0.863 | -0.338 | 0.097 | -0.063 | 1.0 | | | | | | | | |
| $q_{F_1}$ | 0.201 | 0.373 | -0.141 | -0.420 | 0.030 | 0.169 | 0.238 | 1.0 | | | | | | | |
| $q_{F_2}$ | 0.207 | 0.294 | -0.065 | -0.341 | 0.108 | 0.190 | 0.140 | 0.605 | 1.0 | | | | | | |
| $\delta_F$ | 0.153 | 0.242 | 0.207 | -0.262 | 0.053 | 0.207 | -0.222 | 0.449 | 0.514 | 1.0 | | | | | |
| $b_{T_1}$ | 0.101 | -0.016 | -0.023 | -0.010 | -0.573 | 0.257 | 0.0538 | 0.031 | -0.046 | -0.102 | 1.0 | | | | |
| $\delta_T$ | 0.027 | 0.084 | 0.029 | -0.043 | 0.134 | -0.089 | -0.002 | 0.193 | 0.123 | 0.035 | -0.064 | 1.0 | | | |
| $a$ | -0.008 | -0.063 | -0.042 | 0.054 | -0.018 | -0.004 | 0.084 | -0.031 | 0.025 | -0.052 | 0.025 | -0.068 | 1.0 | | |
| $e$ | 0.092 | -0.0036 | -0.133 | -0.042 | 0.079 | -0.389 | 0.119 | 0.001 | 0.003 | -0.041 | -0.231 | 0.084 | 0.024 | 1.0 | |
| $\delta_W$ | 0.046 | 0.066 | -0.019 | -0.089 | 0.146 | -0.315 | 0.026 | 0.141 | 0.146 | 0.197 | -0.336 | 0.237 | 0.004 | 0.833 | 1.0 |

**Table S2** Pearson Rank Correlation coefficient values of young model parameter values generated from bootstrapping.
